# Morphometrics as a conservation tool for rapid in-situ distinction of native and introduced iguanas: comparing *Iguana delicatissima* and *Iguana iguana*

**DOI:** 10.1101/2024.06.10.598237

**Authors:** Matthijs P. van den Burg, Jeroen Kappelhof, Adam Mitchell, Adolphe O. Debrot

## Abstract

Invasive alien species severely impact native and endemic species, disproportionately affecting insular species, and this is especially true for Caribbean reptiles. The Lesser Antillean iguana (*Iguana delicatissima*) experienced a drastic range decline strongly driven by hybridization with non-native green iguanas (NNGI, *Iguana iguana* species complex). With numerous NNGI populations within the Lesser Antilles, it is a matter of time before these arrive on the last *I. delicatisisma* inhabited islands, whereupon rapid in-situ identification of non-native and hybrid animals is essential. However, only a few scale and coloration characters allow in-situ identification of NNGI, which are compromised by introgression. Here we assessed the differentiating power of an additional 20 scale and length-dependent characteristics between the *I. delicatissima* population on St. Eustatius and the established NNGI population on St. Maarten, the main source of arriving stowaway iguanas on St. Eustatius. We identified 14 length-dependent characteristics that significantly differ between *I. delicatissima* and NNGI’s, with multi-variate analysis showing clear morphospace separation and high assignment accuracy of predictive models (>91%). Additionally, the number of femorals and supra digital scales significantly differs between these iguanas. Morphospace knowledge of St. Eustatius’ *I. delicatissima* now allows rapid identification of any divergent iguanas using easy to obtain measurements and meristics. Given intraspecific variation in *I. delicatissima* and green iguanas, we recommend that these characteristics are assessed and validated for other populations, including hybrid individuals. Our work demonstrates the urgent need to invest in baseline morphometric datasets to aid rapid in-situ conservation efforts once NNGI arrive and start to hybridize with *I. delicatissima*.

## Introduction

Invasive alien species are among the most severe actors currently adversely impacting global biodiversity (e.g., Butchart et al., 2010). Insular species are often disproportionately adversely affected by the impacts of invasive species (Dulloo et al., 2002) and with the higher levels of mutualism found in insular ecosystems, impacts can be wide-ranging (Aslan et al., 2013). Besides processes of direct impact such as predation, also processes like invasive hybridization (between native and non-native species) can also cause major declines and extinctions of local endemic populations (Rhymer et al., 1996; Allendorf et al., 2001; Muhfield et al., 2009). Protecting native species from invasive hybridization is essential, especially for rare and endangered species that often occur on islands. However this can be problematic since parental species and their hybrids can be morphologically very difficult to distinguish (e.g., Holsbeek et al., 2008). Morphological identification can be further obscured when hybrids are fertile and introgression between (>)F1 hybrids and parental species occurs (e.g., Haynes et al., 2011), and the presence of this displacing process can go undetected (Lehtinen et al., 2016). As cryptic invasions are common (Morais & Reichard, 2018), local stakeholders are tasked with quick detection of non-native presence and subsequent invasive hybridization to optimize effective species protection.

Within the Caribbean Lesser Antilles, native *Iguana* species are threatened by both invasive hybridization and subsequent competitive replacement with non-native green iguanas from the *Iguana iguana* complex (Breuil et al., 2019; Knapp et al., 2021; van den Burg et al., 2023). This is especially true for the Lesser Antillean iguana (*Iguana delicatissima*), a sister species to *I. iguana*, that has seen its historical range from Anguilla to Martinique decrease by >80% through a multitude of (anthropogenic) factors (Knapp et al., 2014). In the absence of mitigation, its range is projected to decrease to only 1% of its Pre-Colombian range and it is therefore considered as Critically Endangered by IUCN criteria (van den Burg et al., 2018a). Currently, *I. delicatissima* is confined to only a few populations, which either remain pure and inhabit small islets of <2 km^2^ or remain present on larger islands were hybridization has or is occurring (Vuillaume et al., 2015; Angin, 2017; van den Burg et al., 2018b; Pounder et al., 2020). Stakeholders throughout the region are trying to conserve these last remaining populations, a process that is extremely challenging given the number of regionally established non-native green iguana populations that have differing, but only marginally studied (though see Vuillaume et al., 2015; van den Burg et al., 2018b; Pounder et al., 2020) geographic origins from throughout the highly variable *I. iguana* complex (Stephen et al. 2013; van den Burg et al., 2021).

Invasive hybridization with *I. delicatissima* commenced when non-native iguanas arrived in the Lesser Antilles during the 19^th^ century (Breuil, 2013). However, it was not until the 1990s–2000s that this process’ alarming impact on the conservation status of *I. delicatissima* became clear (Lazell, 1973; Breuil et al., 1994; Day & Thorpe, 1996; Day et al., 2000; Breuil, 2000). In fact, Day & Thorpe (1996) already suggested that multi-variate analyses combining morphometrics, and color and scale patterns can quickly differentiate between *I. delicatissima* and *I. iguana*, but these data and results were never published and are believed lost (Roger Thorpe, pers. comm.) Later, Breuil (2013) provided the only published morphological assessment between *I. delicatissima* and green iguanas focussing on scale morphology and coloration patterns. However, subsequent genetic work indicated that introgression can compromise the *in-situ* utility of morphological characters that are currently used to identify non-native iguanas (Vuillaume et al., 2015). Probably, a larger amount of differentiating morphological characters could further improve non-native identification, however since Day and Thorpe (1996), no study has reassessed size-dependent characters to aid differentiation of *I. delicatissima* from invasive green iguanas, even though a recent study did demonstrate that size-dependent characters can aid non-native iguana identification within other native *Iguana* populations in the Lesser Antilles (van den Burg et al., 2023). Here we build on the proposal of van den Burg et al. (2023) to build morphometric baseline datasets of remaining native *Iguana* populations in the Lesser Antilles, with the intended goal of developing this methodology as part of the conservation toolbox for more-effective protection of *I. delicatissima*. We do so by means of an exploratory assessment of variation in size-dependent as well as scalation characters between an *I. delicatissima* population and a regional established non-native green iguana population.

## Materials and Methods

### Study system

Sint Eustatius remains one of the few islands with native *Iguana delicatissima*, however non-native green iguanas occasionally arrive with incoming cargo shipments, presumably from neighboring St. Maarten/Martin (van den Burg et al., 2018b; Debrot et al., 2022). There is a single known case of hybridization between a pure green iguana and *Iguana delicatissima* on St. Eustatius from 2015. That parental non-native female was caught the following year within an ongoing monitoring program and just prior to laying another clutch (Debrot et al., 2022). That program has since resulted in the capture of 10 hybrid iguanas (last in 2020) and additional green iguanas that arrived at the harbor, as recently as March 2024 (Debrot et al. 2022, STENAPA unpublished data). Currently, only a single fully non-native iguana is known to occur on St. Eustatius, which appears to have arrived early 2023 and remains to be captured despite hundreds of search hours. In short, the largely arrested process of hybridization on St. Eustatius stands in high contrast to other islands (Knapp et al., 2014; Vuillaume et al., 2015; Pounder et al., 2020), with any currently present hybrid being the result of a 2023–2024 clutch of the recently arrived non-native iguana in question.

### Fieldwork

We collected data during fieldwork periods on St. Eustatius (2021–2024) and St. Maarten (2024), where we caught iguanas either by hand or using a noose and rope. Photographs were taken of all sides of the head, the anterior dorsal side of the body, the 4^th^ toe of the hind claws, femoral pores, and mid-body ventral scales. Size measures taken following van den Burg et al. (2023), as well as additional characteristics were: snout-vent length (SVL), tail length (TL), lengths of the upper and lower front (UF+LF) and hindleg (UH+LH), length of the 4^th^ toe (TL4), head width (HW), head length (HL), head depth (HD), snout length (SL), eye length (EL), mouth length (ML), tympanum height (TH) and width (TW), and mid-body spine length (MSL). Data on jaw-to-ear (JE) and skull height (SH) were also collected from 2023 onwards (Supplementary info/Appendix 1). After data collection, all iguanas were released in healthy condition at their capture locations. Several characteristics were scored using the photographs: the highest number of femorals on either hindleg (FP), number of supra digital scales on the 4^th^ toe (SD), and the number of ventral mid-body scales within 1 square centimetre (VS). All data and measurements were collected by the first author, and unless specified otherwise, data for bilateral characteristics were collected from the right side (van den Burg et al., 2023). See supplementary information for descriptions of how all measurements were taken.

### Data analyses

Apart from jaw-to-ear height and skull height, all other length variables are already known to be size-dependent and are free of allometry (van den Burg et al., 2023). We first affirmed this for these two new variables. Prior to data analyses all variables were checked for normality and equality of variance. Size-dependent variables were then regressed against SVL for all iguanas of >20 cm SVL, representing late subadults to young and older adults. For each variable we used the resulting SVL-corrected residuals to assess differences between sexes (separately per island) and between *I. delicatissima* vs. non-native iguanas, using either a two-way ANOVA or Kruskal-Wallis test dependent on normality of data distribution and homogeneity of variances. To test the differentiating power of multiple variables, we implemented multi-variate and cluster methodology using principal component (PCA) and discriminant analyses (LDA), which we performed for a dataset using all measured length-dependent variables (except jaw-to-ear and skull height given the later 2023 initiation of inclusion of these variables). We used nine different model-training percentages within our LDA analyses, between 0.50–0.90. A subsequent significance test for these datasets was then performed using a multivariate analysis of variance (MANOVA) for native vs. non-native status. Differences between *I. delicatissima* and non-native iguanas for both meristic variables (femorals and supra digital scales) were assessed using pairwise *t*-tests or Wilcox-Rank Sum tests depending on normality of data distribution and homogeneity of variances. All analyses were done using RStudio version 2023.12.1+402 (RStudio Team, 2019) and R version 4.3.2 (R Core Team, 2023).

## Results

We captured a total of 54 (37 female, 17 male) and 51 (23 female, 28 male) iguanas with a SVL over 20 cm on St. Eustatius and St. Maarten, respectively. An additional 13 smaller iguanas were captured on St. Eustatius, for which we only assessed the meristic characters to those of larger animals (see below). All length variables, including the ventral scale density, were size-dependent for both the St. Eustatius *I. delicatissima* and St. Maarten green iguana population (Table 1). For *I. delicatissima*, size-corrected residuals for each variable, except for head width, eye length and ventral scale density, significantly differed between sexes (Table 1). This was equal for head width and eye length for the green iguana population on St. Maarten, as well as for the size-corrected residuals for head depth and tympanum width (Table 1). Individual comparisons between *I. delicatissima* from St. Eustatius and green iguanas from St. Maarten indicated that 12 SVL-dependent characters significantly differed between these groups. *Iguana delicatissima* had significantly larger upper and lower front legs, a wider head, larger skull height, and higher ventral scale density (Table 1). *Iguana iguana* from St. Maarten had a significantly larger head, snout, and mouth length, larger mid-body spine and tail (Fig. 1), a wider tympanum, and larger head depth (Table 1).

**Table 1.** Morphometric analyses results for *Iguana delicatissima* from St. Eustatius and non-native green iguanas from St. Maarten. R^2^-values of snout-vent length regressions, as well as results of two-sample *t*-tests for size-corrected residual comparisons between the sexes. Significant *t*-test results are indicated by an asterisk for *P* < 0.05. *Iguana delicatissima* and green iguana comparison indicated highlighted in footnote.

| Variable | *Iguana delicatissima* (St. Eustatius) |  |  |  | Green iguanas (St. Maarten) |  |  |  |
| --- | --- | --- | --- | --- | --- | --- | --- | --- |
| | $R^2$ | $t$-value | $df$ | $P$ | $R^2$ | $t$-value | $df$ | $P$ |
| Upper frontleg length<sup>a</sup> | 0.90 | -1.118889 | 53 | 1.782e-06* | 0.92 | -3.8535 | 49 | 0.0003389* |
| Lower frontleg length<sup>a</sup> | 0.94 | -5.2634 | 53 | 2.61e-06* | 0.94 | -3.7883 | 49 | 0.0004153* |
| Upper hindleg length | 0.89 | -4.7913 | 53 | 1.378e-05* | 0.92 | -3.9833 | 49 | 0.0002251* |
| Lower hindleg length | 0.82 | -5.8716 | 53 | 2.896e-07* | 0.91 | -5.394 | 49 | 1.978e-06* |
| Length of 4th toe | 0.62 | -.4.3473 | 53 | 6.281e-05* | 0.82 | -2.1246 | 48 | 0.03879* |
| Head width<sup>a</sup> | 0.90 | 0.42975 | 53 | 0.6691 | 0.89 | -0.058889 | 49 | 0.9533 |
| Head length<sup>a</sup> | 0.84 | -10.208 | 53 | 4.073e-14* | 0.92 | -5.9104 | 49 | 3.215e-07* |
| Snout length<sup>a</sup> | 0.87 | -3.9788 | 53 | 0.0002116* | 0.92 | -2.2601 | 49 | 0.02829* |
| Eye length | 0.76 | -1.622 | 53 | 0.1107 | 0.76 | 0.2433 | 49 | 0.8088 |
| Mouth length<sup>a</sup> | 0.89 | -4.7511 | 53 | 1.584e-05* | 0.94 | -3.6358 | 49 | 0.0006642* |
| Head depth<sup>a</sup> | 0.75 | -4.2645 | 52 | 8.474e-05* | 0.91 | -0.97638 | 49 | 0.3337 |
| Tympanum height | 0.75 | -7.4614 | 53 | 8.164e-10* | 0.78 | -2.7732 | 49 | 0.007829* |
| Tympanum width<sup>a</sup> | 0.81 | -3.4082 | 53 | 0.001256* | 0.74 | -1.8725 | 49 | 0.06711 |
| Mid-body spine length<sup>a</sup> | 0.57 | 31.672<sup>b</sup> | 1<sup>b</sup> | 1.825e-08<sup>b*</sup> | 0.71 | -5.5733 | 49 | 1.056e-06* |
| Jaw height | 0.65 | -8.4164 | 38 | 3.266e-10* | 0.82 | -7.457 | 49 | 1.304e-09* |
| Skull height<sup>a</sup> | 0.72 | -9.6515 | 38 | 9.065e-12* | 0.83 | -7.346 | 49 | 1.934e-09* |
| Ventral scale density<sup>a</sup> | 0.82 | -0.67037 | 48 | 0.5058 | 0.76 | -2.0728 | 43 | 0.04421* |
| Tail length<sup>a</sup> | 0.79 | -5.2758 | 39 | 5.228e-06* | 0.84 | -2.715 | 20 | 0.01314* |

<sup>a</sup> $P < 0.05$ for size-corrected comparison between *I. delicatissima* and green iguanas.
<sup>b</sup> Results of Krus-Wall chisquared test due to not normal and unequal variance.

**Figure 1.**
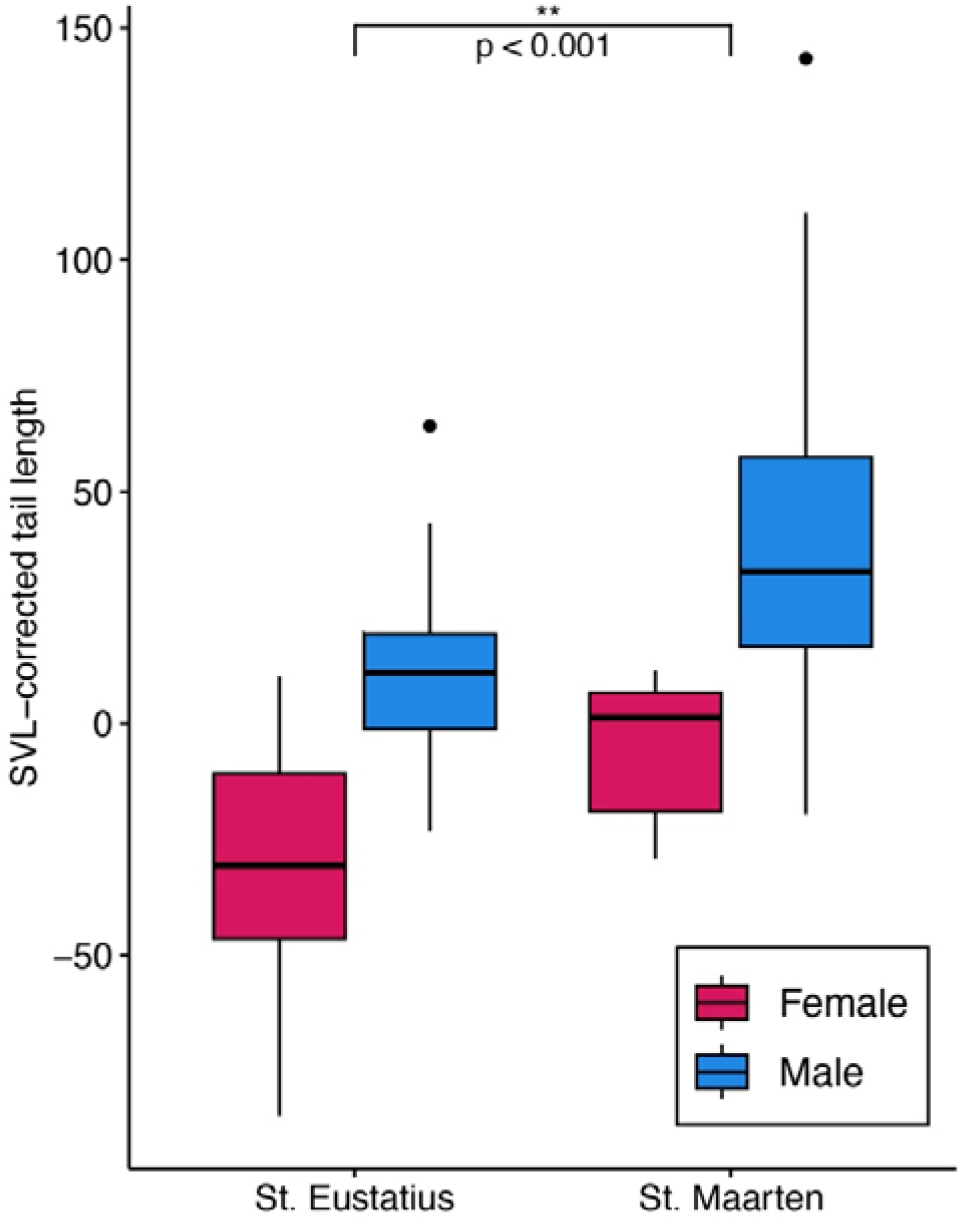
Differences in tail length between between sexes and *Iguana* populations on St. Eustatius and St. Maarten non-native and native iguanas on Saba. Data are SVL-corrected residuals from iguanas of >20 cm SVL.

The number of supra digitals on the 4^th^ toe differed significantly between sexes for *I. delicatissima* (*W* = 152.5, *p* < 0.001), but not for green iguanas from St. Maarten (*W* = 265.5, *p* = 0.2812). The number of supra digitals was significantly higher for green iguanas than for *I. delicatissima* (*t* = −12.915, *df* = 106, *p* < 2.2e-16; Fig. 2B). The number of femoral pores was not significantly different between sexes, but *I. delicatissima* had significantly more femorals compared to green iguanas from St. Maarten (*W* = 3051, *p* < 2.2e-16; Fig. 2A). Femoral and supra digital counts for small-sized *I. delicatissima* fell within the range of those of larger *I. delicatissima*; *n* = 13, supra digitals: 30–35, femorals: 20–22.

**Figure 2.**
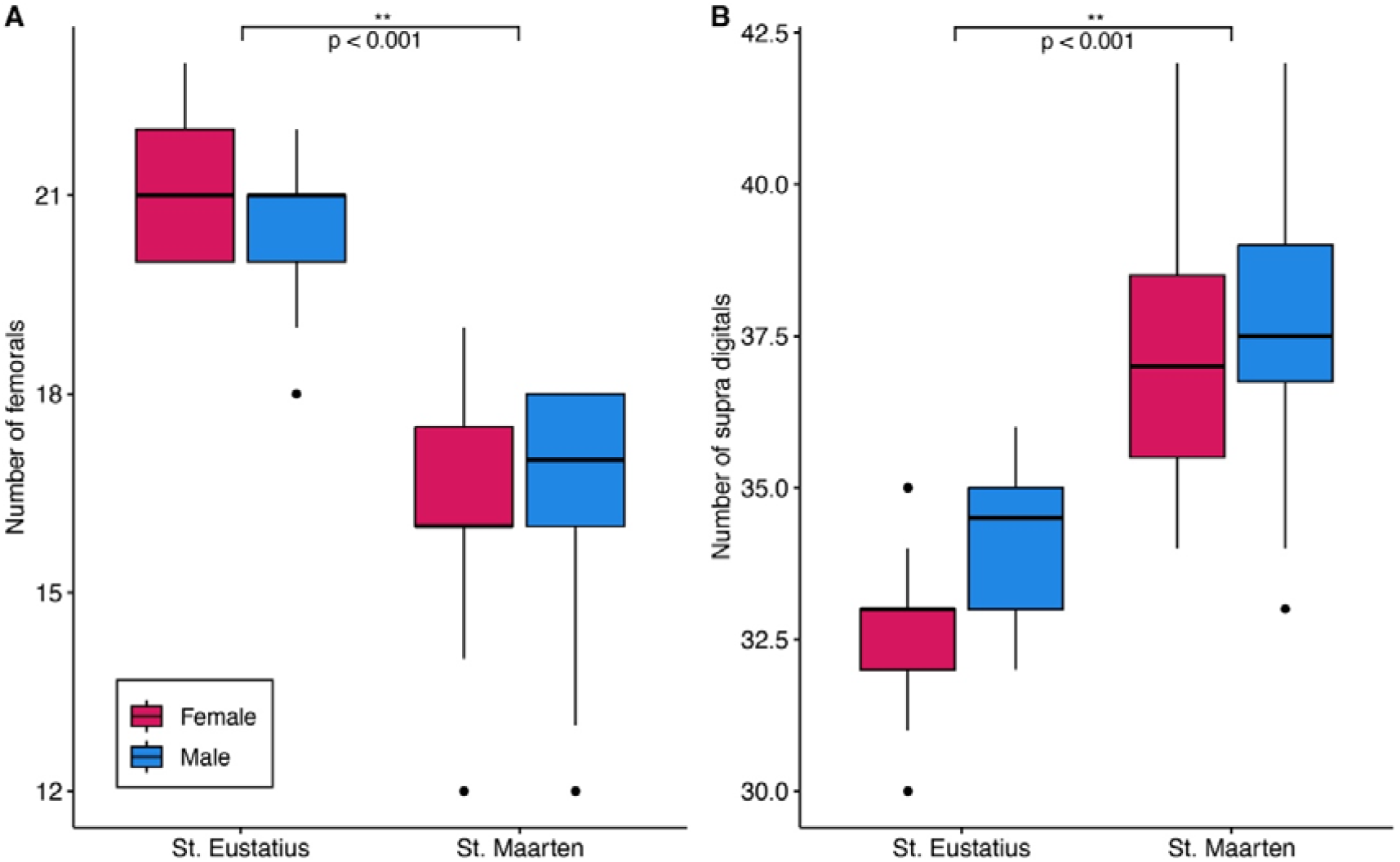
Variation in (A) femorals and (B) supra digitals on the 4^th^ toe between sexes and *Iguana* populations on St. Eustatius and St. Maarten.

The PCA analysis for all size-dependent variables (except jaw-to-ear and skull height) shows a near absence in morphospace overlap between *I. delicatissima* from St. Eustatius and green iguanas from St. Maarten (Fig. 3). PC1-axis contributes most to separation between sexes, and PC2 between *I. delicatissima* and green iguanas. Highest loadings for PC1 were UH, LH, HL, SL, ML, and MSL, and for PC2 UF, LF, HW, SL, HD and MSL. The nine LDA models using 50-90% training input had high accuracy (94–100%) for correctly assigning *I. delicatissima* and green iguanas. The one-way MANOVA for island status was significant (*F*_1,91_ = 30.605, *p* < 0.001, Wilk’s Λ = 0.154). These analysis steps were repeated with all variables with significant difference being found between the islands: UF, LF, HW, HL, SL, ML, TW, SH, MSL, TL. Given a high percentage of individuals had regenerated tails (Table 1), fewer individuals had no missing data: 25 for *I. delicatissima*, and 19 for green iguanas from St. Maarten.

**Figure 3.**
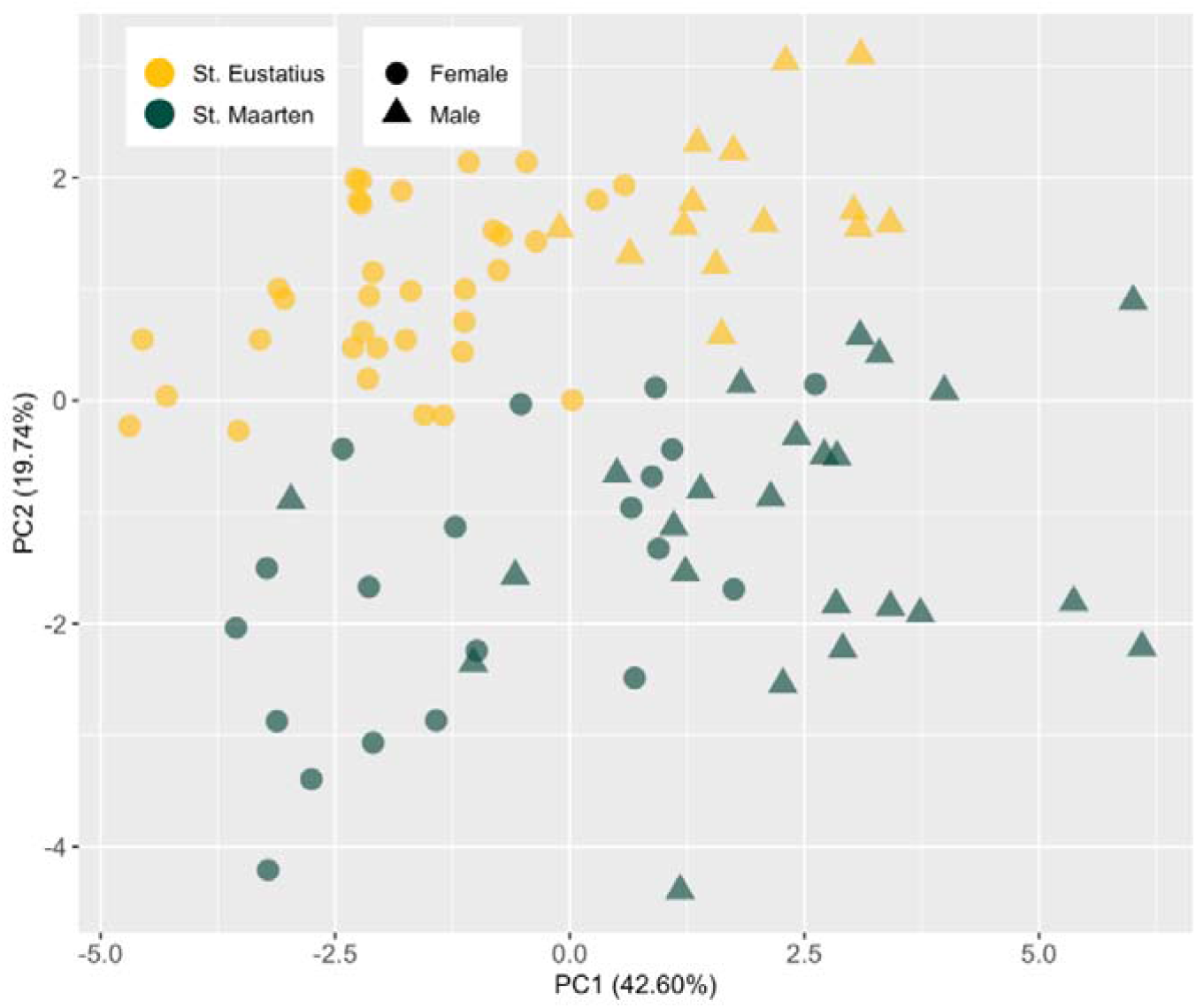
Principal component analysis plot summarizing variation across residuals of all SVL-dependent variables in morphospace, except jaw-to-ear and skull height.

## Discussion

Our assessment highlights numerous morphometric and meristic characters that allow rapid *in-situ* identification of non-native green iguanas that can be used to boost on-ground conservation management of the remaining *Iguana delicatissima* populations. Although this study is restricted to a comparison between only two populations, our results develop an important new tool to help protect the St. Eustatius *I. delicatissima* population from its most important regional source of invading green iguanas, St. Maarten. While Day & Thorpe (1996) indicate the presence of intraspecific variation in *I. delicatissima* that appears associated with differences in climate and island banks, our effort should be viewed as a time-saving exploratory example for our regional partners, to help them develop their own island-specific assessments. Islands where hybridization is ongoing allow assessing at which extent and degree of introgression these characters are compromised, and which combination of characters maximizes correct distinction of *I. delicatissima* within a continuous introgression scenario.

Since Breuil (2013), no additional differentiating morphological assessments between *I. delicatissima* and green iguanas have been published, despite the knowledge that morphometric differences exist (Day & Thorpe, 1996). Here we identified an additional set of 14 characters that significantly differed between *I. delicatissima* from St. Eustatius and green iguanas from St. Maarten (Table 1; Fig. 3). Since we focused our explorative effort on larger iguanas, follow-up assessments should investigate the utility of the identified 12 size-dependent characters for smaller-sized (<20cm SVL) native and non-native individuals. Scale counts, other than size-dependent characters, are less likely to have ontogenetic variation and we identified two useful meristic characters that were also used by Lazell (1973); the max number of femorals (Fig. 2A), and number of supra digitals on the 4^th^ toe (Fig. 2B). Data from some (*n* = 13) smaller iguanas indeed show concordance with the variation found for larger individuals. Therefore, we urge more attention towards identification of additional meristic characters with differentiating power between native vs. non-native iguanas. Although Lazell (1973) contrarily did not find a difference between *I. delicatissima* and *I. iguana* for these two meristic characters, his results were likely impacted given the inclusion of introgressed populations from the Guadeloupe archipelago (Breuil, 2013). Lastly, here we provide a preliminary comparison of total length between *I. delicatissima* and regionally established non-native green iguanas, as an important dominance feature in iguanas (Dugan, 1982). The displacement of *I. delicatissima* by non-native green iguanas is well known, although the mechanisms underlying this process are yet to receive research attention. So far aggressive outcompetition including greater strength of green iguanas has been proposed (Knapp et al., 2014; Vuillaume et al., 2015), and a singular record of hybrid fecundity suggests these might have higher fitness compared to pure *I. delicatissima* (van Wagensveld & van den Burg, 2018). Here we provide a first data-driven insight on male-to-male dominance between green iguanas and *I. delicatissima*, since our results show that tail size is larger for green iguanas for equal SVL (Fig. 1). Green iguanas thus attain a relatively larger lateral profile compared to *I. delicatissima*. We note this assessment only included half of our dataset due to the large number of individuals with a partially regenerated tail (see *df* statistic in Table 1). We thus recommend future studies to identify a proxy for total tail length, to allow inclusion of all animals in data analyses and usage within multi-variate analyses. Circumference of the tail base at the cloaca, or at a SVL-dependent distance from the cloaca would be first options to explore.

Compared to *I. delicatissima*, intraspecific variation is more evident within green iguanas (Day & Thorpe, 1996; Breuil, 2013; Stephen et al., 2013; van den Burg et al., 2021; van den Burg et al., in prep.), although a full range-wide assessment has not been performed. Established non-native *Iguana* populations within the Lesser Antilles have different, and often multiple, geographic origins (Vuillaume et al., 2015; van den Burg et al., 2018; Pounder et al., 2020; De Jesús Villanueva et al., 2021; Breuil et al., in prep.). Although larger non-native sample sizes (>20) have only been analysed from Anguilla, Antigua, Guadeloupe and St. Maarten/Martin (Vuillaume et al., 2015; Pounder et al., 2020; Breuil et al., in prep.; van den Burg et al., in prep.), some populations still await assessment e.g., Barbuda and Dominica. The St. Maarten/Martin population currently has the most complex introgressive history, with green iguanas from multiple major mtDNA clades hybridizing as well as evidence of some remaining *I. delicatissima* introgression (Breuil et al., in prep.). This high variation from mixed origins could explain some observed contrast between dimorphic differences concerning three variables compared to *I. delicatissima* (Table 1). Our results indicate that, head depth and tympanum width show dimorphic differences in *I. delicatissima* but not in green iguanas, even though, this was reversed for ventral scale density. In reptiles, the density of ventral scales has been linked to climatic variation, including for *Anolis* (Thorpe, 2017), and preliminary data from the *Iguana iguana* complex suggests a potentially similar relationship (van den Burg et al., in prep). Hybridization with or among non-native species occurs in numerous systems and provides opportunities to study potential selection and adaptive introgression (Whitney & Gabler, 2008; Schulte et al., 2012; Chaplin et al., 2022; DeVos et al., 2023). This is especially because the range of green iguanas includes multiple geographic and climatic regions and the species’ multi-origin populations present throughout the Caribbean could provide insights into these processes.

Population-specific baseline datasets can aid rapid in-situ identification of non-native iguanas and could replace the long process of genetic identification when adequate differentiating characters and datasets are available (van den Burg et al., 2023). However, fieldwork duration and data collection are restrained by limited time and financial resources and ideally each data-collected character has some degree of differentiating power between native and non-native individuals. Our preliminary assessment aids regional conservation partners in prioritizing characters to measure, especially if limited resources do not allow the recommended complete local assessments. Whilst size-dependent characters can only be measured in-situ, photographing meristic characters can provide an opportunity for subsequent *ex-situ* data collection. We used this combined *in-/ex-situ* workflow to assess the number of femorals and supra digitals, and ventral scale density. In conclusion and adding to Breuil (2013), we here identify several morphological characters that can be used to aid the rapid distinction of *I. delicatissima* from non-native iguanas allowing better and more timely protection of the last remaining *Iguana delicatissima* populations from the threat of invasive hybridization. With the native morphospace of the St. Eustatius *I. delicatissima* population defined, rapid identification of any divergent (hybrid or introgressed) iguana should be more straightforward using a number of measurements and meristics. We call on conservation and governmental bodies to provide local conservation organizations with rapid funding to construct local population-specific baseline datasets. For islands with ongoing introgression (St. Barthelemy, Martinique, La Desirade and Dominica), financial aid should also include funding for an initial genetic assessment.

## Acknowledgements

We are grateful for financial support that allowed us to perform this study from Rotterdam Zoo, and the Ministry of Agriculture Nature and Food Quality (LNV) through the Wageningen University BO research program (BO-43-117-006) under Wageningen University and Research project number 4318100346-1. We thank the St. Eustatius National Parks Foundation and Nature Foundation St. Maarten for supporting this project, which approved and authorized our fieldwork methodology. Data were collected on St. Eustatius within the iguana monitoring program of the St. Eustatius National Parks Foundation.

